# The ecological memory of landscape complexity shapes diversity of freshwater communities

**DOI:** 10.1101/2024.11.01.621539

**Authors:** Andrea Tabi, Edineusa Pereira Santos, Gabriel Brejão, Tadeu Siqueira

## Abstract

Land use change affects the biodiversity of terrestrial landscapes as well as of the freshwater systems they surround, and thereby numerous ecosystem services that are essential for human well-being. Previous research suggests that freshwater systems show delayed responses to abrupt disturbances such as deforestation, however less is known about whether there is a legacy effect of landscape history on freshwater communities under a long history of land use change. We addressed this research gap here by first quantifying the historical complexity of landscapes, including natural and agricultural formations, in terms of their composition and configuration in 26 tropical catchment areas surrounding 101 streams over a 30-year period. We identified clear evidence for a memory effect of past events, whereby historical landscape complexity, measured as the information entropy of landscape composition, positively affected freshwater biodiversity. Finally, using a causal discovery approach, we showed that species richness was causally related to landscape complexity only when historical values were incorporated. Our results corroborate previous work on the positive effect of landscape complexity on biodiversity, and also confirm the role of historical contingencies in predicting future ecological outcomes.

## 1 Introduction

Land use change has become one of the major causes of biodiversity losses around the world (38, 41). Human activities have been transforming landscapes for thousands of years, destroying natural habitats and causing species extinctions (19). The historical dependency of biodiversity on land use change has been demonstrated in many systems (33, 47), including forests (43) affected by various human activities (8, 23, 29, 53). These ecosystem responses to past events and conditions — also called the ecological memory — are crucial for increasing future resistance and resilience of ecological systems (3).

Previous empirical studies on ecological memory have mainly focused on the effects of abrupt conversion of one natural land cover type (e.g., natural forest) to one anthropogenic land use type (e.g., pasture) (6, 24, 52). However, most landscapes have a long history of land use change and exist as complex mosaics comprising both natural and modified formations. Capturing the complexity of such mixed landscapes is not an easy task because biological and physical properties and processes in a landscape interact simultaneously between its parts (28). Recent studies have measured landscape complexity as the percentage of natural elements in the landscape (7), using metrics based on Shannon information entropy (40), or various metrics to measure spatial configuration of land cover (10, 40, 62). Overall, these studies indicate that higher landscape complexity leads to higher biodiversity in agricultural landscapes (20) and to an increased provision of ecosystem services such as water quality, pollination, pest regulation, carbon storage (16), and crop production (40). However, less is known about to what degree the historical complexity of landscapes shape biodiversity in areas with a long history of land use change.

The conversion of natural land cover to intensive land uses, such as agriculture, forestry, and pasture, affects both terrestrial and riverine communities at different spatial and temporal scales (1, 2). That is because the hierarchical structure of riverine networks, which are embedded in terrestrial drainage basins, make them both receivers and transmitters of materials and organisms. They receive sediment, nutrients, contaminants, and organisms from the surrounding landscape and from upstream; they process and transmit these downstream (17). In riverine communities, invertebrates include species that are sensitive to environmental change and that play pivotal, diverse roles in food web dynamics. While random demographic events and density independent and dependent processes shape the species composition of aquatic invertebrates at the local scale, dispersal plays a major role in distributing individuals among localities within the catchment area (37, 57). Dispersal of riverine invertebrates occur via two main pathways: the aquatic stages disperse through watercourses, while winged adults disperse along and across riparian buffers (32). Thus, the structural complexity of the surrounding landscape, including its composition and spatial configuration, should play a major role in the assembly of riverine biodiversity.

Ecological memory has been previously quantified using weight functions, which determine the functional form of discounting past values, such as hyperbolic or exponential decay. For instance, the hyperbolic and exponential decay functional form has been used in psychology to model forgetting — known as Jost’s Second Law of Forgetting (31, 50)—, and in economics, to explain future consumer choices (36, 56). Other more flexible frameworks such as Bayesian hierarchical models (30), stochastic antecedent modelling (44), or random forests (4), estimate the relative importance (weight) of the past time points of the explanatory variables on the target variable with predefined time lags. In this study, we present a nonlinear, nonparametric framework to quantify discrete memory effects of landscape complexity on freshwater biodiversity using observational data on 101 tropical freshwater communities across 26 river catchments in Brazil, combined with annual historical land use cover data between 1985 and 2015. By using this approach, we aimed to identify past events with most impact on the current diversity without having a predefined time lag or weight function. Furthermore, we present a causal discovery analysis to map the causal links between landscape complexity metrics, diversity and other biotic and abiotic factors that potentially affect diversity.

## 2 Methods

### 2.1 Study sites and data

We compiled landscape data by analyzing maps of land use and land cover from the Corumbatí River basin, state of São Paulo, southeastern Brazil (Fig.1). This basin comprises an area of 1,700 km^2^, and its natural vegetation cover was originally dominated by semi-deciduous forests and with patches of Savanna (51). Initial deforestation in the basin began in the early 18th century with farming, followed by an expansion of pasture land and crop land, mainly sugar cane, which modified the natural vegetation cover to small fragments (21). More recently, the amount of native forest cover has been increasing due to natural vegetation recovery in abandoned agricultural lands (21, 39). Within the Corumbatí River basin, we delineated 26 catchments using a SWAT model (Soil and Water Assessment Tool), based on a Digital Elevation Model (DEM) that was obtained from SRTM (Shuttle Radar Topography Mission, 30 m of spatial resolution). We used MapBiomas 8.1 with 30 m of spatial resolution (55). We quantified the land use and land cover classes (in m^2^) yearly between 1985 and 2015.

**Figure 1.**
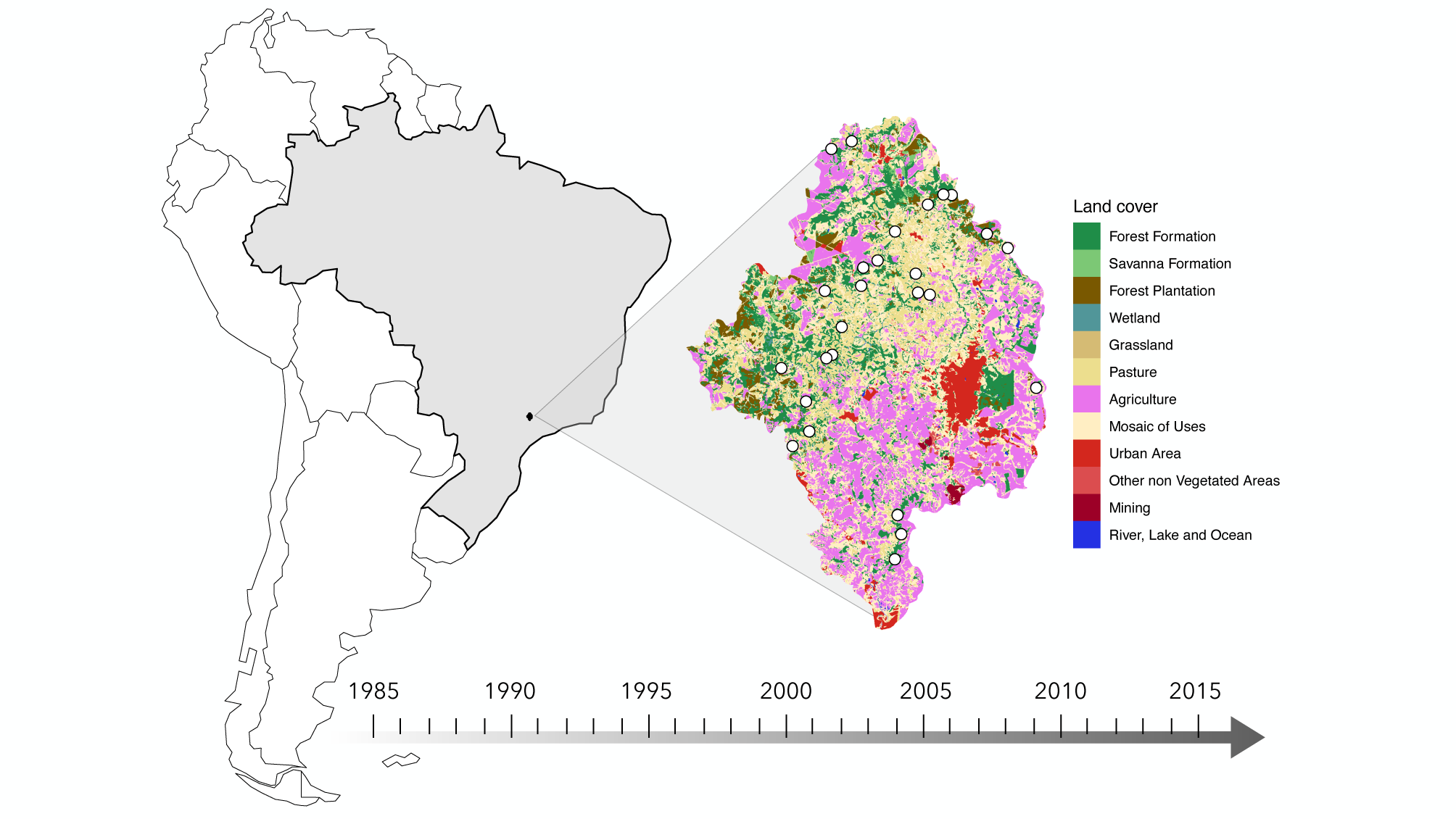
Land cover data and sampling sites. Our study site is the Corumbatí River basin, state of São Paulo, southeastern Brazil. We collected landscape data by analysing maps of land use and land cover from 26 catchments between 1985 and 2015, where we surveyed 101 streams for aquatic insects in 2015.

We surveyed 101 streams distributed within 26 catchments, in which we collected immature aquatic insects predominantly in the dry season between May and August 2015. We used a kick-net, with 1 *m*^2^ area and 250 *µm* mesh, to collect a generous sample in each stream by gently kicking the stony substrate of four riffles for a total of two minutes (30 seconds each). In each of the four riffles, we sampled aquatic insects. The identification of the collected specimens to species or morphotype was based on taxonomic keys and original descriptions (14, 15). To estimate body size, we averaged the size of the three largest specimens of each species. Feeding strategy of species was collected from literature (9, 48, 58). In each stream, we also obtained measurements of turbidity, conductivity and dissolved oxygen with a multiparameter YSI 556 MPS probe (YSI Inc., Ohio). Stream width, water temperature and flow velocity were measured as well. In each stream, we took water samples that were analysed for total nitrogen and total phosphorus, following national standards for Brazil (27). We also visually estimated the proportion of organic detritus (small litter patches) intercepted by rocks in the stream riffles and the proportion of shading.

### 2.2 Measuring landscape complexity and memory effects

We used three entropy metrics to quantify landscape complexity of each catchment area. First, we calculated the normalised Shannon information entropy to calculate compositional complexity of the different land cover and land use types in each catchment area: 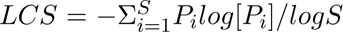, where *P_i_*is the relative area of cover type *i*. The values of landscape composition vary from 0 to 1, with zero representing landscapes dominated by a single land cover type (low entropy) and with 1 representing landscapes with equally distributed land cover types. Then, we used two spatial entropy metrics, the Wasserstein distance (61, 62) and the Boltzmann approach, to capture the configuration of the landscape mosaic (10, 12). The Wasserstein metric calculates *W_c_*, the distance between frequency of states (land cover types) and a reference distribution (Dirac delta distribution) and *W_s_*, the distance between the repetition of spatial configuration and reference state (Dirac delta distribution); *LCW* = (1 − *W_c_*) · (1 − *W_s_*). Both *W_c_* and *W_s_* are normalized between [0,1] by the number of pixels (62). The Boltzmann entropy approach for landscape mosaic calculates the total length along edges of pixels where two different cover types touch. First, a landscape mosaic is spatially permuted 5000 times, producing a distribution of randomized landscape mosaic patterns. Then, the total edge length is calculated for the landscape and for the permuted patterns representing the possible microstates. The entropy (*S_B_*) is calculated from the fitted log-normal distribution of microstates (12). Finally, we calculated the normalised Boltzmann entropy as 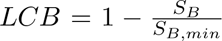, where *S_B,min_* is the entropy of the theoretical minimal edge length (54). We also quantified the co-occurrence of the agricultural land and forest cover as the relative number of neighbouring cells of each cover type.

In order to detect the memory effect of landscape complexity on species richness in riverine communities, we used a Stochastic Local Search (SLS) algorithm called Threshold Accepting (TA) (26). The memory weights were constrained as 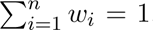, where *n* is the number of time points. The algorithm maximized the Spearman’s correlation coefficient between diversity and the sum of the weighted landscape complexity time series 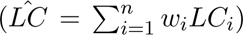. The advantage of stochastic optimization techniques over other similar algorithms, such as simulated annealing (SA), is that depending on the choice of the threshold it does not get stuck in local minima due to its different acceptance rules (18). TA accepts every new configuration as long as it is not much worse than the old one, whereas SA accepts worse solutions with very small probabilities. On average, TA provides better solutions with shorter computational time compared to SA. We ran the algorithm 500 times for each case in order to obtain a distribution of correlation coefficients. Each time the algorithm was run for 5 · 10^5^ iterations. Then, we calculated the average weight values and confidence intervals.

### 2.3 Discovery of causal structure

In the causal discovery analysis we included the measured species richness in 101 streams, community-weighted species traits (predation and body size), locally measured variables (organic detritus in stream, shading, temperature and stream width), and the three landscape complexity metrics. First, we found the skeleton (i.e. undirected causal links among the variables), then oriented edges. We used the IC algorithm (45, 46) to construct the skeleton. The algorithm takes as input a stable probability distribution generated by some underlying causal structure and outputs a *pattern* (partially directed graph). To find the skeleton, i.e. undirected causal links, for each pair of variables 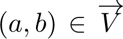 searches for a set *S_ab_* such that (*a* ⊥⊥ *b*|*S_ab_*). Then, an undirected graph *G* is defined such that vertices (*a, b*) are connected with an edge if and only if no set *S_ab_* can be found. The IC algorithm is based on conditional independence testing, in case of continuous variables partial correlation tests. We also tested these models by setting across different significance levels (*α* [0.01, 0.1]) of the conditional independence tests.

## 3 Results

Overall, ten land cover categories including natural and non-natural formations between 1985 and 2015 were identified and measured within the catchment areas of the Corumbatí river basin, south-eastern Brazil (Fig.S1). All catchments had a long history of land use modification to various degrees (Fig.S2). The temporal changes in landscape composition measured as the Shannon information entropy increased during the study period, which was mostly driven by changes in agricultural land cover expansion (Fig2D). Natural forest cover was temporally stable during over the 30-year period. The spatial co-occurrence of agricultural land cover and forest cover was calculated as well, however they were very highly correlated with the land cover values. Changes in landscape configuration were quantified by two spatial entropy metrics; Wasserstein (LCW) and Boltzmann (LCB) entropy. Wasserstein entropy captures the distribution of composition and patch sizes, and was strongly correlated with landscape composition. Boltzmann entropy, which measures the length of the overall perimeter of land type covers, showed no strong trend across the study period (Fig.2C). The two spatial entropy metrics exhibited no association. The analysis showed that landscape composition had the strongest effect on diversity, with the highest average memory weight in 2012 (Fig.3A). Using the concurrent value of landscape composition, the effect is significantly lower (*ρ* = 0.22) compared to incorporating the effect of historical values. Similarly, landscape configuration calculated as the Wasserstein metric, showed the importance of 2012, however its overall effect on biodiversity was lower than landscape composition (Fig.3B). The effect of landscape configuration calculated as the Boltzmann metric remained very low after the optimisation (Fig.3C). We also analysed the effect of agricultural land cover on biodiversity, however we did not recover memory effect of the year 2012 (Fig.S4). The concurrent correlation value between agricultural cover and diversity was higher than the optimised values (Fig.3D).

**Figure 2.**
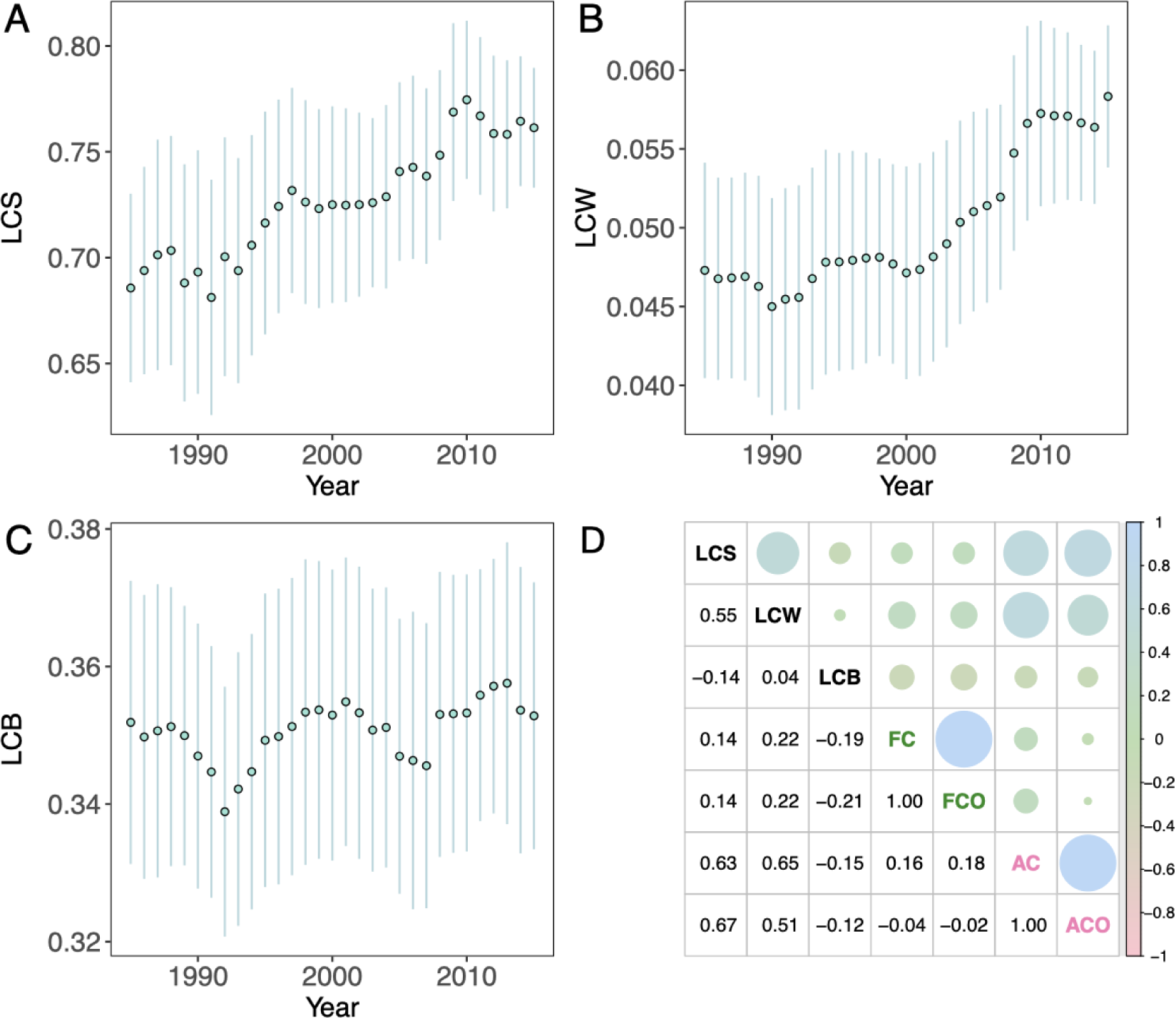
Measuring landscape complexity in river catchments. (**A-C**) Time series of landscape complexity metrics (Shannon (LCS), Wasserstein (LCW) and Boltzmann (LCB) entropy). calculated annually over a 30-year period in each catchment (colors). (**D**) shows the Spearman’s correlation between landscape complexity metrics and agricultural cover (AC) and co-occurrence (ACO) and forest cover (FC) and co-occurrence (FCO) time series.

**Figure 3.**
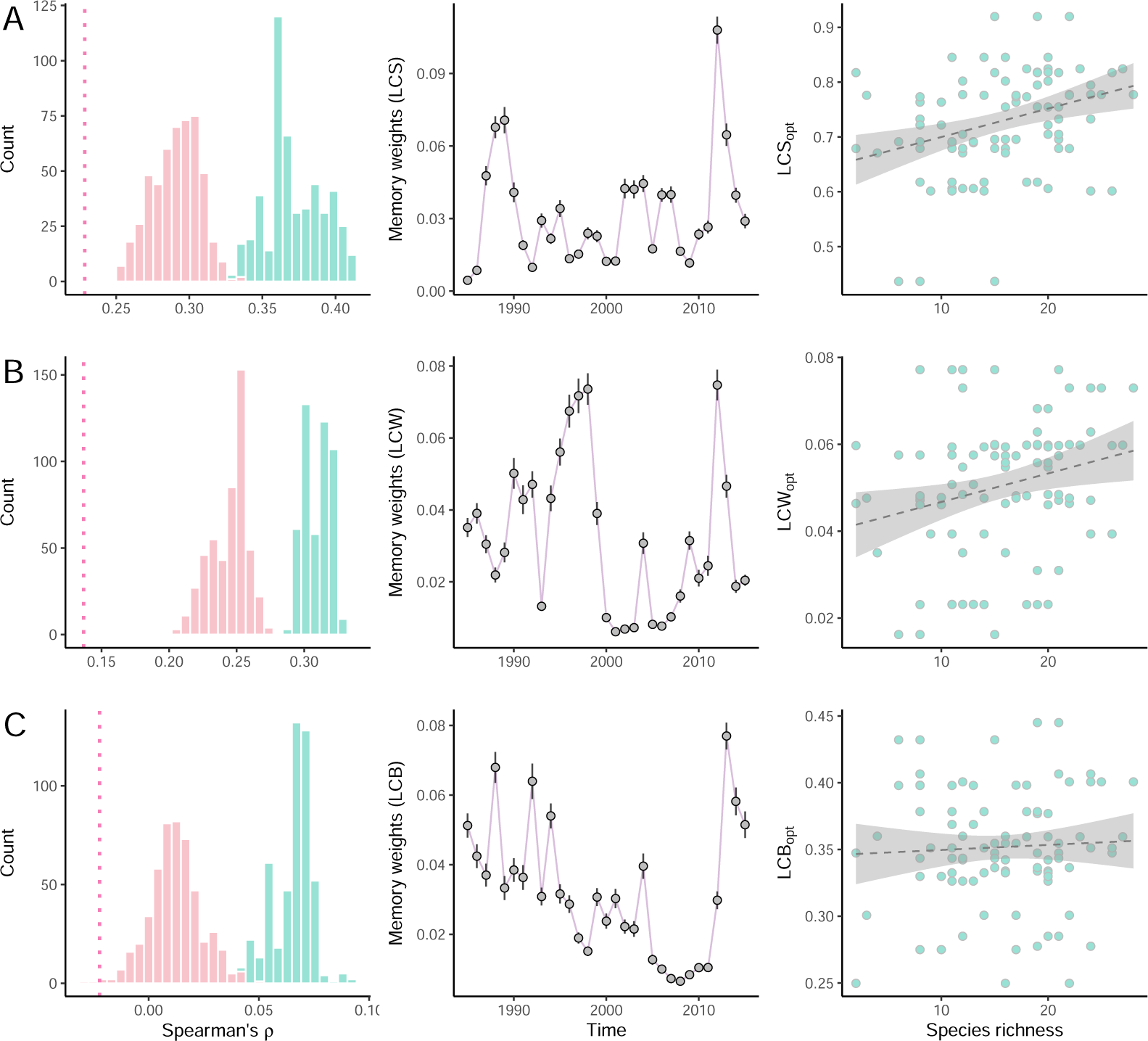
Estimated average memory weights. Memory weights were estimated using a stochastic optimisation approach for the Spearman’s *ρ* between diversity and the three landscape complexity metrics. On the left hand side, the distribution of the values with optimised (blue) and values with random memory weights (pink) of Spearman’s *ρ* are depicted. The vertical dotted line shows the concurrent correlation value. In the middle, the ensemble average of memory weights by year are shown. On the right hand side, the relationship is plotted between the optimised landscape entropy metrics and species richness.

Observational data are naturally confounded. So, in order to establish causality between species richness and landscape complexity, and to confirm a true memory effect, we conducted a causal inference analysis. For this analysis, in addition to the three landscape complexity metrics, we included community-weighted traits (predation and body size) and site-specific variables (organic detritus in stream, shading, water temperature and stream width) (Fig.S5). We tested the causal graph at different significance levels with and without incorporating the optimised historical landscape complexity metrics. Then, we identified the threshold at which the focus link between species richness and landscape complexity appeared in the causal structure. When we estimated the causal structure with concurrent values (Fig.4A), landscape complexity did not have a causal effect on diversity or any other variables. However, the inclusion of historical landscape metrics resulted in a different causal structure. We found that historical landscape composition was the only landscape metric that directly affected species richness (Fig.4B). Higher values of landscape composition, representing more evenly distributed land cover classes, increased species richness. The only abiotic variable that directly influenced species richness was the amount of organic detritus, whereby higher amounts of organic detritus in streams contributed to higher diversity. Predation, as the most important biotic factor in our analysis, decreased species richness, whereby predators had larger average body size.

**Figure 4.**
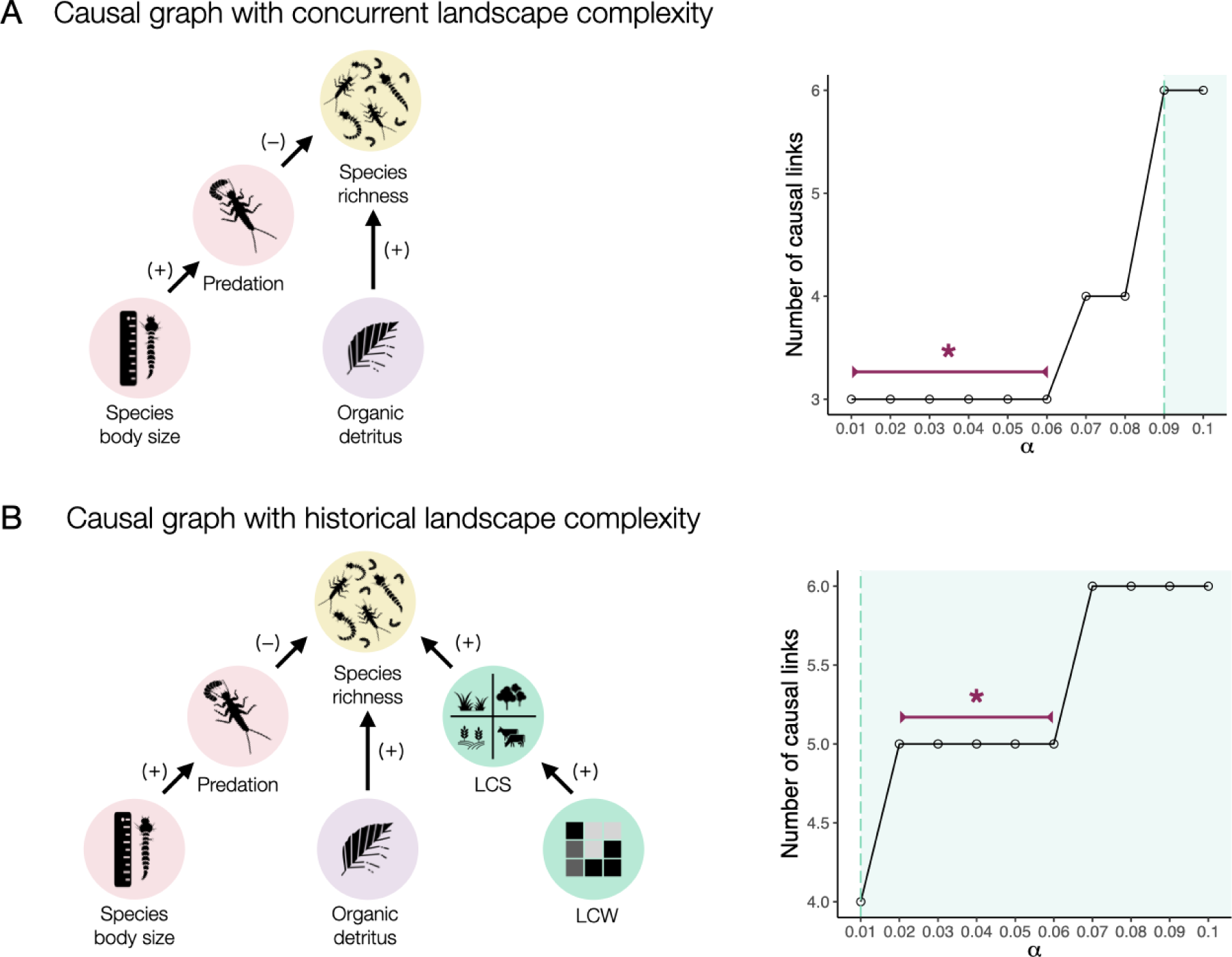
Causal structure. (A) The number of causal links resulted from the causal discovery analysis with concurrent landscape entropy values increased with the significance level (*α*) of conditional independence tests, where the green dashed line indicates the threshold, where the link between species richness and landscape composition first appeared. (B) The causal structure including historical landscape compelxity metrics shows that species richness of freshwater communities is directly affected by historical landscape composition, predation and organic detritus in stream.

## 4 Discussion

Our findings confirmed the presence of memory effects of historical landscape complexity on the biodiversity of tropical riverine systems. More specifically, historical landscape composition, measured as Shannon information entropy, had a strong influence on species richness. Furthermore, we identified the most relevant point in time when compositional change in land cover affected species richness. The analysis revealed that agricultural land expansion drove changes in landscape composition on average, however, the importance of historical agricultural land cover did not peak in the same years as the historical landscape composition. The other major changes in many locations around this time period can be attributed to the increase in forest plantations (Supplementary Information Fig. S2), a pattern that mirrors observations for the entire Atlantic Forest Biome. This simultaneous land cover change over time highlights the need for landscape entropy metrics that are able to capture the complex nature of such structural changes in landscapes. Structural changes can happen gradually or abruptly over time, which is reflected in the ecological memory of the community. This means that nonparametric approaches are the most suitable for detection of memory in ecological system(4, 44), opposed to continuous decay functions (34).

Our analysis identified 2012 as the most critical point in time when compositional changes in land cover affected species richness. The landscape changes around this period primarily reflected an increase in agricultural cover. This shift can be partially attributed to the introduction of the Native Vegetation Protection Law (NVPL, Law #12,651/2012) in Brazil. Despite the establishment of innovative control programs and incentives to promote compliance with the NVPL, the new law reduced restoration requirements, allowed certain forms of agriculture in areas designated for native vegetation, and granted amnesty for fines to rural landowners and landholders not complying with previous regulations (5). Given that some of these changes were anticipated prior to the law’s implementation, it is reasonable to conclude that the landscape alterations observed around 2012 left a legacy effect on species richness. For instance, Nunes et al. (42) found higher deforestation rates within riparian zones compared to non-riparian areas in 2012. Given the critical role of riparian forests in buffering changes within catchment areas for aquatic communities, even minor forest losses in these zones can have substantial impacts on freshwater biodiversity (13).

Capturing the heterogeneity of land cover types has been a primary challenge in understanding the interactions between landscapes and underlying ecological processes. The use of different entropy metrics to describe the complexity of landscape mosaics, however, has been deeply rooted in landscape ecology (11, 59). The lack of importance of spatial configuration as a driver of species richness suggests that landscapes are highly structured, rendering configurational information that is redundant in explaining biodiversity patterns. This result suggests that the applied spatial entropy metrics did not capture the most relevant feature of spatial heterogeneity with regard to species richness, as it has been shown before (40).

Our causal analysis confirmed that landscape composition was not only associated but also causally related to species richness, and most importantly not confounded by any other factors. Species richness was also directly related to local biotic and abiotic factors, such as predation and amount of organic detritus in streams, respectively. In line with previous studies (57), predation had a negative effect on species richness, a relationship that has been suggested to be particularly stronger in the tropics (25). Body size, which is considered a master trait that scales with many other species traits and interactions (35), was causally-related to predation, whereby predators tended to have larger body sizes (60). A higher accumulation of organic detritus in streams contributed to higher species richness. A major proportion of the organic detritus in streams comes from terrestrial areas (22). However, landscape complexity metrics were not causally connected to species traits, nor to the amount of organic detritus in streams. Because landscape complexity metrics were measured within a 30-meter radius, we may have underestimated the role of riparian buffers of native vegetation, which, even when narrow, are the major contributors of organic matter inputs into streams (49).

Our research demonstrated that discrete historical events of landscape change impose constraints on freshwater biodiversity. This agrees with previous work on biological legacies as part of the ecological memory of communities (3). We also confirmed that composition is an essential feature of landscapes to explain species richness in tropical riverine systems. Following a causal inference approach, we showed how landscape complexity and other biotic and abiotic factors influence local freshwater biodiversity. Although more research is needed to explore the connection between biodiversity and landscape complexity and how it varies across different scales and contexts, our research indicates a strong role of landscape-level historical contingencies in predicting freshwater biodiversity.

## Data accessibility

The data supporting the results will be archived on data depository upon acceptance of the manuscript.

## Acknowledgements

The acquisition of data used in this study was supported by grant #2013/50424-1, São Paulo Research Foundation (FAPESP). TS was supported by grant #2021/00619-7, FAPESP. This study was financed in part by the Coordenação de Aperfeiçoamento de Pessoal de Nível Superior - Brasil (CAPES) - Finance Code 001. We thank Andŕe M. de Roos for the insightful comments on the manuscript.

## Author contribution

AT designed and implemented the analysis and wrote the manuscript. EPS and GB collected the data. All authors interpreted the results and edited the manuscript.

## Supporting Information

**Figure S1.**
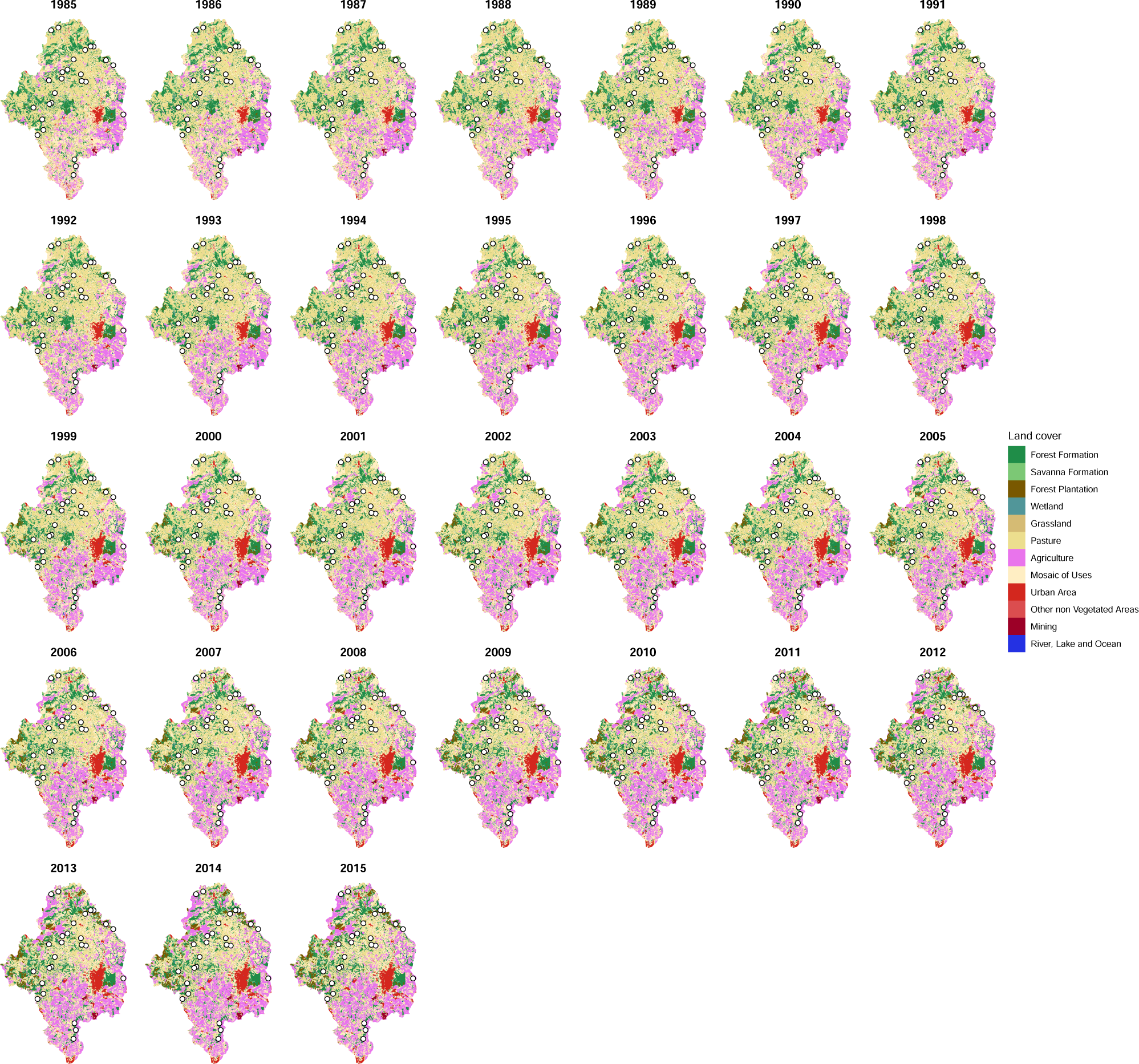
Regional land cover 1985-2015.

**Figure S2.**
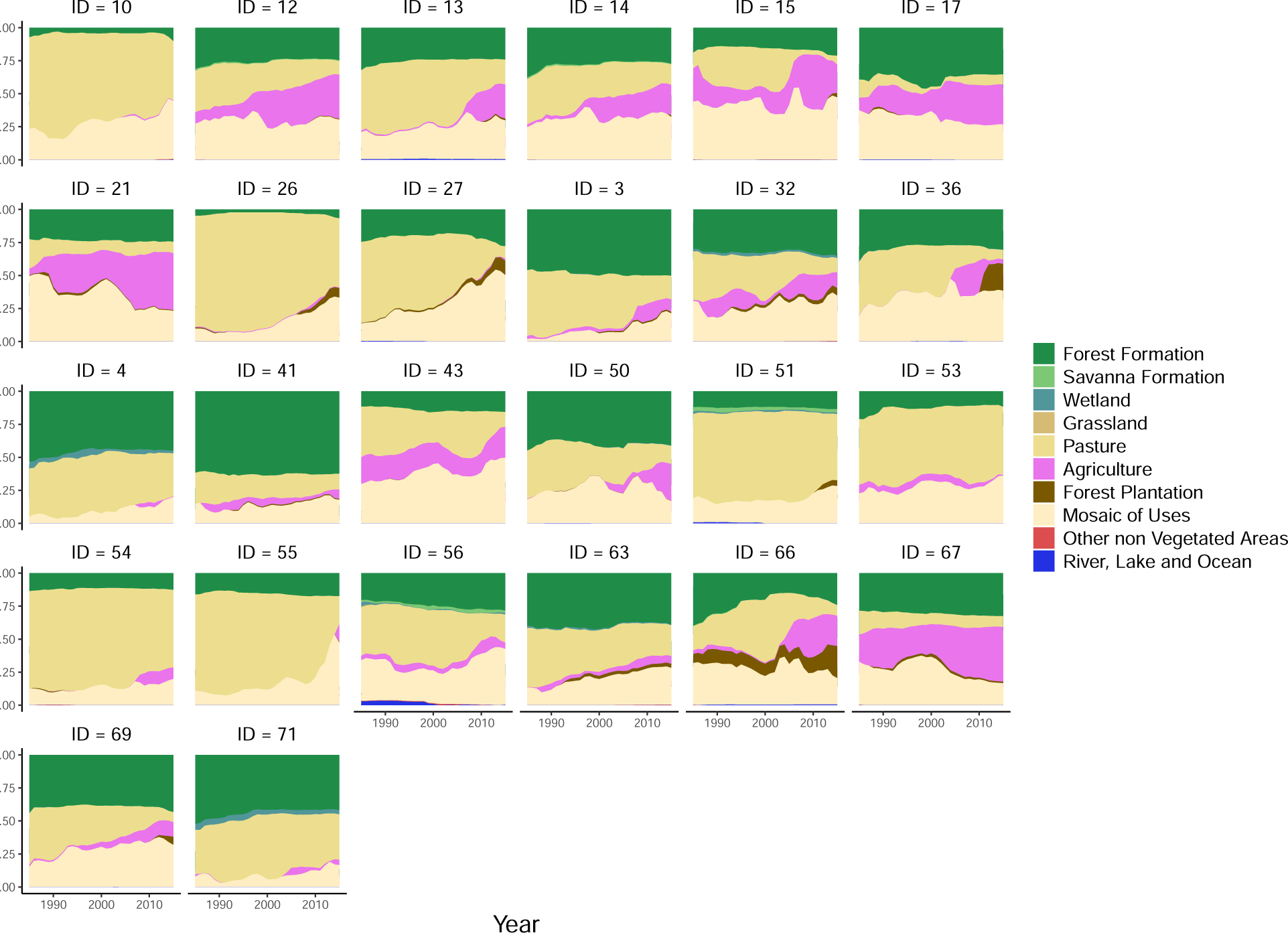
Catchment land cover composition 1985-2015.

### Parametric estimation of memory effects with weight functions

We applied four decay functions (*D*(*t, δ*)) to landscape entropy time series; binary step-based, exponential, hyperbolic and linear decay functions with decay rates (*δ*) between [0, 1] (Fig.S3) calculated as follows:

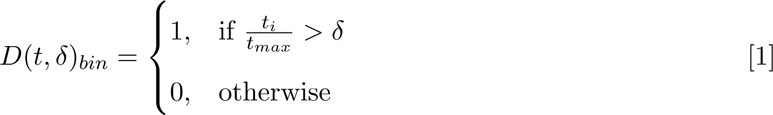

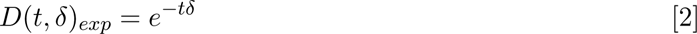

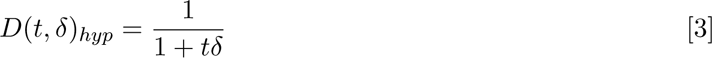

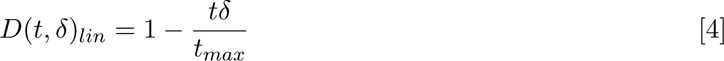

Then, we calculated the Spearman’s correlation between the average value of the weighted landscape entropy time series and species richness across decay rates. The memory effect can be detected as changes in correlation coefficients, e.g. we expect a decrease in correlation with higher decay rates if memory effect is present.

**Figure S3.**
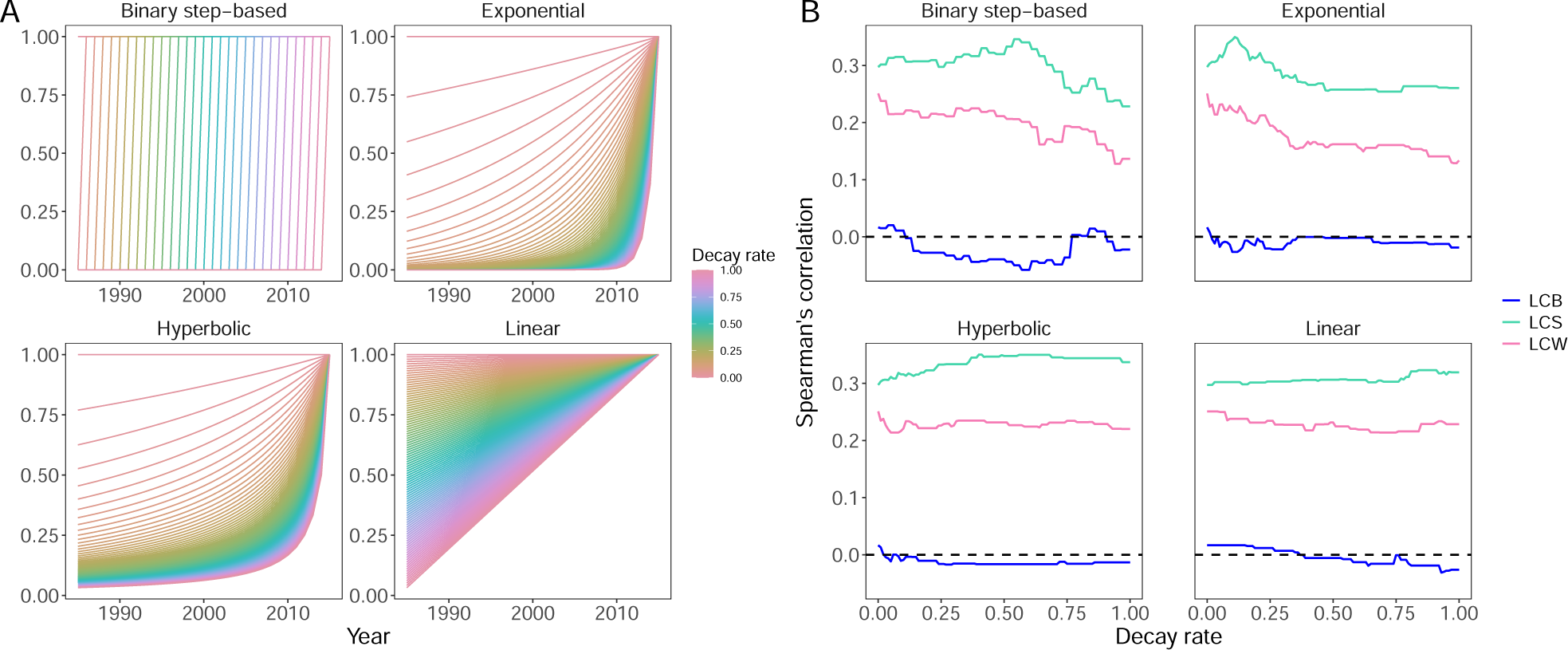
Measuring memory effects. (A) Four different decay functions were applied to landscape entropy time series. (B) The Spearman’s correlation coefficient between species richness and landscape complexity by decay rates was studied in order to identify memory effects. The correlation between landscape composition and richness decreased with increasing decay rates when binary step-based and exponential functions applied, whereas hyperbolic decay seem to capture the functional form of memory effect, reaching the highest Spearman’s coefficient. The memory effect of landscape composition was revealed by the decline in the relationship between species richness and averaged discounted landscape complexity as decay rates increased. This analysis showed that higher decay rates in exponential and binary step-based functions applied to landscape composition weakened its effect on species richness, whereas in the case of hyperbolic functional form, higher decay rates slightly improved the correlation. This result means that the removal or quick discounting of historical values of landscape composition reduced the predictive power of historical landscape composition on species richness.

**Figure S4.**
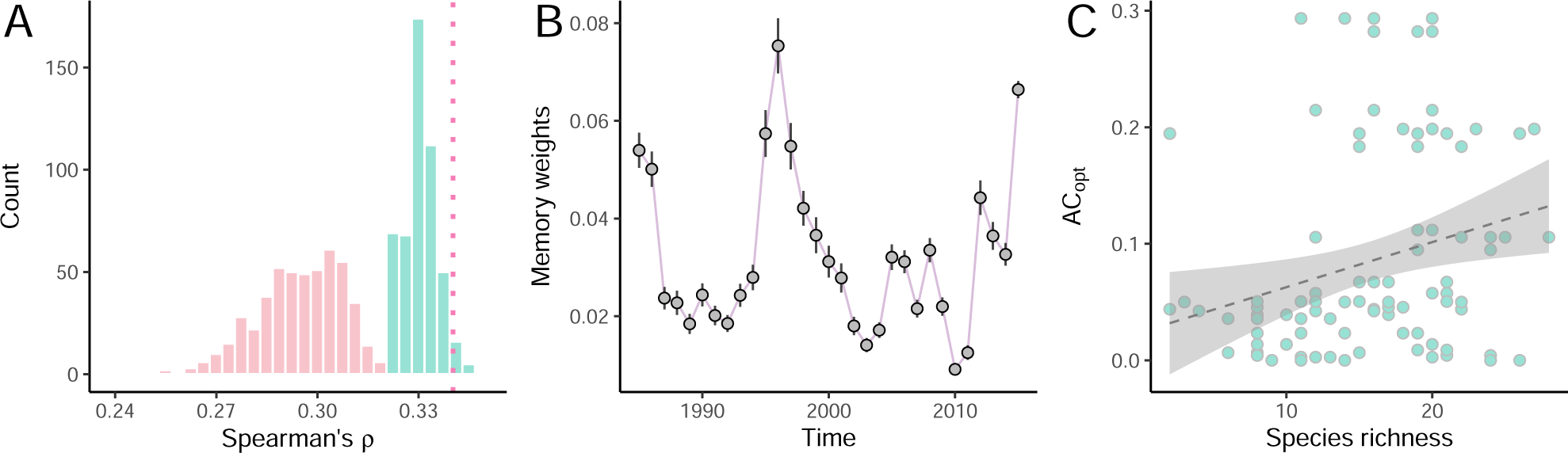
Estimated average memory effects of agricultural cover.

**Figure S5.**
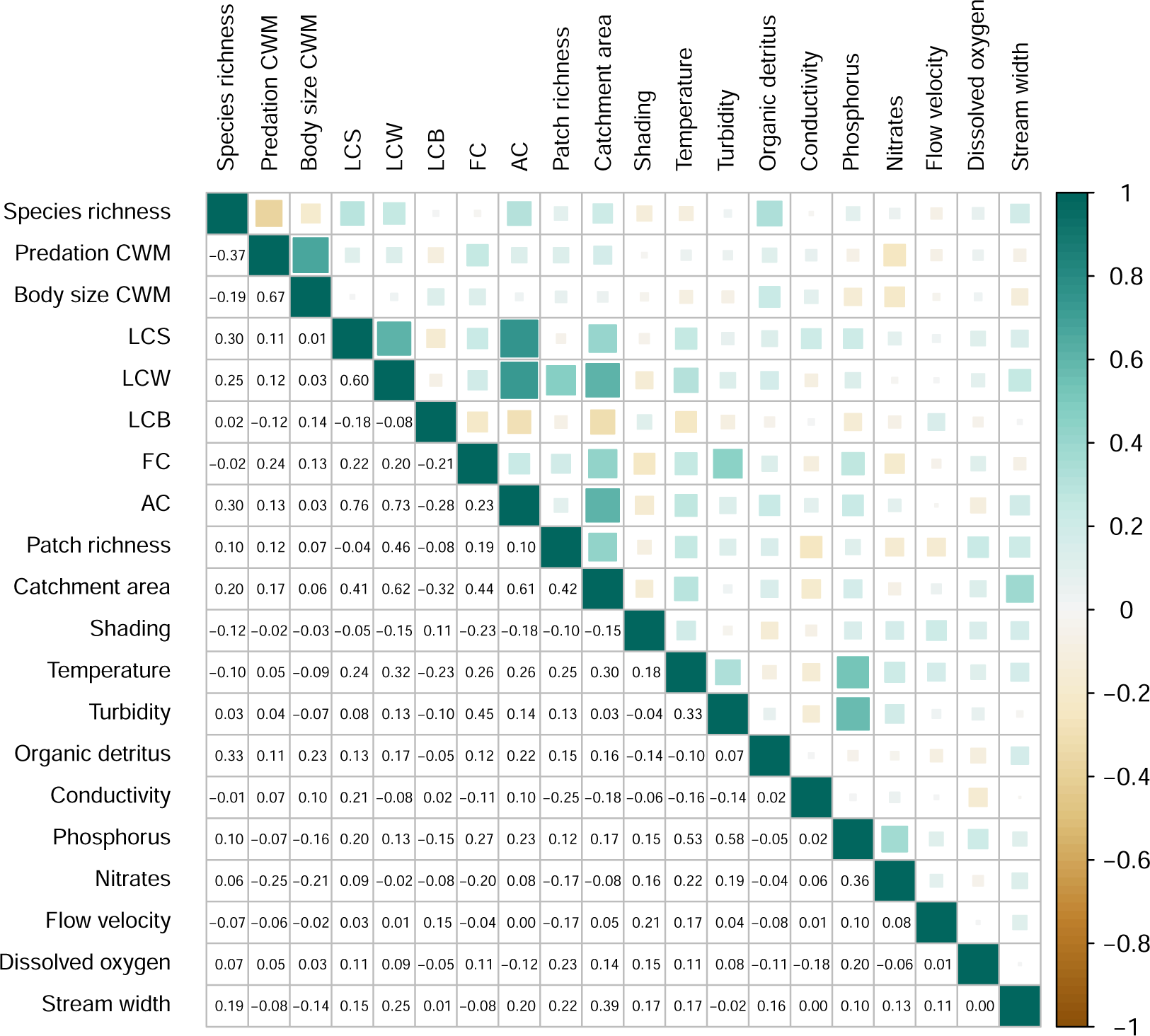
Spearman’s correlations among variables.

